# Revisiting criteria for plant miRNA annotation in the era of big data

**DOI:** 10.1101/213314

**Authors:** Michael J. Axtell, Blake C. Meyers

## Abstract

MicroRNAs (miRNAs) are ~21 nucleotide-long regulatory RNAs that arise from endonucleolytic processing of hairpin precursors. Many function as essential post-transcriptional regulators of target mRNAs and long non-coding RNAs. Alongside miRNAs, plants also produce large numbers of short interfering RNAs (siRNAs), which are distinguished from miRNAs primarily by their biogenesis (typically processed from long double-stranded RNA instead of single-stranded hairpins) and functions (typically via roles in transcriptional regulation instead of post-transcriptional regulation). Next-generation DNA sequencing methods have yielded extensive datasets of plant small RNAs, resulting in many miRNA annotations, occasionally inaccurately curated. The sheer number of endogenous siRNAs compared to miRNAs has been a major factor in the erroneous annotation of siRNAs as miRNAs. Here, we provide updated criteria for the confident annotation of plant miRNAs, suitable for the era of “big data” from DNA sequencing. The updated criteria emphasize replication, the minimization of false positives, and they require next-generation sequencing of small RNAs. We argue that improved annotation systems are needed for miRNAs and all other classes of plant small RNAs. Finally, to illustrate the complexities of miRNA and siRNA annotation, we review the evolution and functions of miRNAs and siRNAs in plants.

## Introduction

Small regulatory RNAs are a major feature of eukaryotic transcriptomes. In plants, many such RNAs are produced by Dicer-Like (DCL) proteins via excision from double-stranded RNA (dsRNA) or from the paired regions of single-stranded, stem-loop RNAs. After excision, single-stranded RNAs from the initial duplexes become bound to effector proteins in the Argonaute (AGO) protein family. The assembled AGO-small RNA complex is then able to identify RNA targets based on complementarity between the target RNA and small RNA. Successful target recognition allows the AGO protein to directly or indirectly perform a negative regulatory function.

Plants produce many distinct types of DCL/AGO-associated regulatory small RNAs (Axtell, 2013). Our discussion focuses on three of the major types of plant small RNAs:

1. MicroRNAs (miRNAs). These are defined by the precise excision of the initial duplex from the stem of a stem-loop precursor RNA. The initial duplex consists of the strand that will become the functional miRNA, and a partner that is less frequently bound to an AGO protein, the miRNA* (“microRNA-star”). Plant miRNAs generally target mature mRNAs that are either protein-coding or long, non-coding RNAs (lncRNAs). In plants, RNA targets of miRNAs are repressed, a result of either or both mRNA destabilization and translational repression. Plant miRNAs tend to be 21 or 22 nucleotides (nts) in length.
2. Phased siRNAs (phasiRNAs). PhasiRNAs are products of processive cleavage of double-stranded RNAs (dsRNAs) in regular increments (duplexes) from a well-defined terminus. The terminus is typically defined by a miRNA- or siRNA-directed, AGO-catalyzed cleavage event that occurred on a single-stranded precursor. After cleavage, the precursor is converted to dsRNA by the activity of an RNA-Directed RNA Polymerase (RDR), which in most known cases is an ortholog of *Arabidopsis* RDR6. However, long inverted repeats may also give rise to phasiRNAs, even in the absence of a small RNA trigger - for example, the SRK gene in some Arabidopsis ecotypes (Lu et al., 2006). When there is a single, initiating cleavage event, the dsRNAs all begin at the same nucleotide, so that when the siRNAs are successively processed from that terminus, they are produced in a sequential pattern, termed ‘phasing’. This pattern of phasing can be used to identify secondary siRNAs based on reference-aligned sRNA-seq data. These siRNAs are ‘secondary’ when their biogenesis is dependent on the initial miRNA or siRNA interaction (aka, the “trigger”). Like plant miRNAs, phasiRNAs are frequently 21 nts in length, although one major subgroup is 24 nts in length. Both coding mRNAs and long non-coding RNAs can be phasiRNA sources.
3. Heterochromatic siRNAs (hc-siRNAs). Like phasiRNAs, hc-siRNAs are excised from RDR-dependent dsRNA precursors (typically made by orthologs of *Arabidopsis* RDR2). But since the dsRNA precursors of hc-siRNAs are ~30-50 nts (Blevins et al., 2015; Zhai et al., 2015), these give rise to only one siRNA and hence there is no possible phasing from a single precursor. Their defining features are their origins from intergenic, repetitive, and transposon-related genomic regions, and they are typically 24 nts in length. They function to target nascent, chromatin-tethered non-coding RNAs; target recognition causes *de novo* deposition of DNA methylation on the adjacent genomic DNA, ultimately leading to the reinforcement of heterochromatic histone marks.

Readers interested in the details of biogenesis and molecular functions of miRNAs, phasiRNAs, and hc-siRNAs should consult more comprehensive or specialized reviews (Rogers and Chen, 2013; Fei et al., 2013; Matzke and Mosher, 2014). Here we focus on the new knowledge and challenges that have arisen from the application of high-throughput small RNA sequencing (sRNA-seq) to the discovery and study of these three major classes of plant small RNAs in diverse plants. We also envision this article as an update to a decade-old publication that laid out the criteria for the annotation of plant miRNAs (Meyers et al., 2008), taking into account the many technological and conceptual advances made since that time.

## Update on Criteria for Plant miRNA Annotations: The Urgent Need to Minimize False-Positives

The first coordinated effort to define community standards in miRNA annotations relied on a simple set of criteria involving a combination of evidence of expression (accumulation on an RNA blot or in a cDNA sequencing project) and biogenesis (presence of a hairpin, conservation of the miRNA, or reduction of accumulation in a *dicer* mutant background) (Ambros et al., 2003). This first effort occurred well before the full complexity of endogenous small RNAs was understood, and was later found by many (including us; (Meyers et al., 2008)) to be inadequate, especially in the context of sRNA-rich plants. In particular, the accumulation of a small RNA on an RNA blot merely proves it accumulates, but does not differentiate between miRNAs, endogenous siRNAs, and partially degraded fragments of RNA unrelated to any gene-regulatory mechanism. Conservation is not definitive as a criterion for annotation both because there is no *a priori* reason that a miRNA need be conserved (and indeed, many appear not to be; see below), and because there are other types of small RNAs that are conserved. Reduction in a *dicer* mutant is also not definitive in plants because plants express multiple *DCL* genes with partially redundant roles in both miRNA and siRNA biogenesis (Gasciolli et al., 2005; Rajagopalan et al., 2006), as well as in other events such as tRNA processing (Cole et al., 2009; Martínez et al., 2017). In other words, reduction or loss in a *dcl* mutant proves the RNA is not just a random degradation product unrelated to gene regulation, but does not categorically differentiate between miRNAs and endogenous siRNAs.

Careless application of the 2003 criteria to modern deep sequencing data would have led to a deluge of false-positive miRNA annotations. Realizing this, in 2008, we coordinated the publication of a consensus opinion among many plant small RNA researchers on a better set of criteria for plant miRNA annotations (Meyers et al., 2008). The 2008 criteria essentially stated that the only necessary and sufficient criteria for annotating a miRNA locus is the demonstration of precise processing of a miRNA/miRNA* duplex from a computationally-predicted hairpin RNA precursor that met certain minimal structural criteria (Table 1). One of the major requirements in terms of making a novel annotation is the sequencing of the exact miRNA*. miRNA* abundance may be one or two orders of magnitude less than that of the mature miRNA, and so the requirement of miRNA* sequencing necessitates deep sequencing coverage.

**Table 1.** Updated criteria for plant miRNA annotations

|  | 2008 Criteria | 2018 Criteria |
| --- | --- | --- |
| 1 | One or more miRNA/miRNA* duplexes with 2 nt 3' overhangs | Add requirements that exclude secondary stems or large loops in the miRNA/miRNA* duplex, and limit precursor length to 300 nts. |
| 2 | Confirmation of both the mature miRNA and its miRNA* | Disallow confirmation by blot – sRNA-seq only |
| 3 | miRNA / miRNA* duplex contains $\leq 4$ mismatched bases | Up to 5 mismatched positions, only 3 of which are nucleotides in asymmetric bulges |
| 4 | The duplex has at most one asymmetric bulge containing at most two bulged nucleotides |  |
| 5 | $\geq 75\%$ of reads from exact miRNA or miRNA* | Include one-nt positional variants of miRNA and miRNA* when calculating precision |
| 6 | Replication suggested but not required | Required – novel annotations should meet all criteria in at least two sRNA-seq libraries (biological replicates) |
| 7 | Homologs, orthologs, and paralogs can be annotated without expression data, provided all criteria met for at least one locus in at least one species | Homology-based annotations should be noted as provisional, pending actual fulfillment of all criteria by sRNA-seq |
| 8 | miRNA length not an explicit consideration | No RNAs $< 20$ nt or $> 24$ nts should be annotated as miRNAs. Annotations of 23- or 24-nt miRNAs require extremely strong evidence. |

In our opinion, the 2008 criteria for plant miRNA annotations have held up quite well in the last ten years. However, our own research experiences in the intervening time suggest the following improvements (summarized in Table 1):

1: The hairpin of a miRNA precursor is a defining characteristic, a component of all reliable miRNA-identification algorithms. Using conserved miRNAs as exemplars, there are key features of the foldback or hairpin that are important for Dicer recognition and consistent processing (Cuperus et al., 2011; Bologna et al., 2013b; 2013a; Chorostecki et al., 2017): (i) typically a single miRNA:miRNA* duplex (thus far not more than three duplexes, a number observed in miR319/159); (ii) the region of this foldback that gives rise to the miRNA duplex should not contain secondary stems or large internal loops (larger than five nucleotides) that interrupt the miRNA:miRNA* duplex; (iii) while miRNA foldbacks of plants are often longer than those of animals, most foldbacks are just a few hundred nucleotides. These structural characteristics govern processing including the base-to-loop versus loop-to-base direction of duplex production. Foldbacks longer than 300 nucleotides should not be annotated as miRNA-producing loci.
2: Expression of one or more miRNA/miRNA* duplexes should be observed by high-throughput small RNA sequencing (sRNA-seq), as opposed to reliance on RNA blotting. RNA blotting for small RNAs carries the risk of probe cross-hybridization. sRNA-seq methods are now routine and should be the standard method of expression analysis used to support novel miRNA annotations in plants.
3 and 4: Changes in structural criteria. Up to five mismatched positions between the miRNA and miRNA*, only three of which can be asymmetrically bulged, should be allowed. These changes correspond to observed extremes among validated miRNAs. Note that these criteria were separate in the 2008 ‘rules’ (Table 1), but are similar and thus combined in the 2018 criteria.
5: Precision of miRNA/miRNA* processing and positional variants. The 2008 criteria suggested that 75% or more of all aligned small RNAs at a locus should correspond to the exact miRNA or miRNA* sequences. Our experiences with sRNA-seq data from many species since then indicates this is too strict. In particular, we observe that 5’ and especially 3’ positional variants of miRNAs and miRNA*s are quite common in sRNA-seq data. These variants likely result from a combination of DCL cleavage at different positions (including DCL imprecision, perhaps a characteristic of recently-evolved miRNAs (Bologna et al., 2013b)), post-dicing modifications of the 3’ end (Zhai et al., 2013), and sample degradation. Thus the revised calculation of precision includes the counts of the exact mature miRNA and its corresponding exact miRNA*, as well as all one-nucleotide variants thereof. A one-nucleotide variant is defined as a small RNA for which both the 5’ and 3’ ends are each within one nucleotide of the corresponding ends of the exact miRNA or miRNA*. Thus for each miRNA and miRNA* there are eight possible one-nucleotide variants. Precision is thus re-defined to be the sum of reads for the miRNA, miRNA*, and their respective one-nucleotide variants divided by the total number of reads aligned to the locus. When calculated in this manner, precisions for high-confidence microRNA loci tend to be around 90%; the minimum allowable percentage remains 75%.
6: Replication. Annotations of novel miRNA families require support from independent small RNA-seq libraries. That is, all of the updated criteria must be fulfilled in at least two distinct sRNA-seq libraries (not technical replicates). Ideally, these replicates would come from different tissues, developmental stages, or treatments, as miRNAs show a higher degree of qualitative reproducibility across any of these than siRNAs. However, biological replicates of the same tissue/stage/treatment could suffice for miRNAs for which expression is truly specific to one condition.
7: Annotation by homology. We feel that any annotations based on homology alone to a known miRNA family should be regarded as provisional, and clearly marked so, until such time as all the required sRNA-seq based criteria are met for the newly annotated locus. This will slow down the rate of propagation of errors caused by homology-based annotations of families erroneously annotated in the first place.
8: miRNA length. One effective method to minimize false positive annotations is to categorically disallow annotations of miRNAs that are not the expected size. We are not aware of any evidence demonstrating that miRNAs < 20 nt or > 24 nt are loaded onto any plant AGO proteins. Unless such evidence emerges, RNAs < 20 or > 24 nts in length should never be annotated as miRNAs. hc-siRNAs are mostly 23 or 24 nts in length, and they are extremely numerous in many plant sRNA-seq libraries. In contrast, *bona fide* 23 or 24 nt miRNAs are very rare. The huge number of 23- or 24-nt hc-siRNAs coupled with scant experimental evidence for 23- or 24-nt miRNAs suggests that avoiding annotations of 23- or 24-nt RNAs as miRNAs is prudent. Any exceptions should have truly extraordinary evidence of miRNA biogenesis and function, including precise miRNA/miRNA* accumulation in four or more separate sRNA-seq libraries and ideally a direct demonstration that they post-transcriptionally regulate target RNAs in a miRNA-like manner (i.e. translational repression, target cleavage, and/or phasiRNA accumulation). One potential caveat to this rule is that it remains possible that some plant species generate numerous 23- or 24-nt miRNAs, but these species have not yet been characterized or had extensive small RNA analyses.

Figure 1 shows an example of two *Arabidopsis thaliana* loci analyzed using sRNA-seq data from inflorescences. While the locus ath-*MIR399b* meets all of the revised criteria described here, ath-*MIR405a* does not.

**Figure 1.**
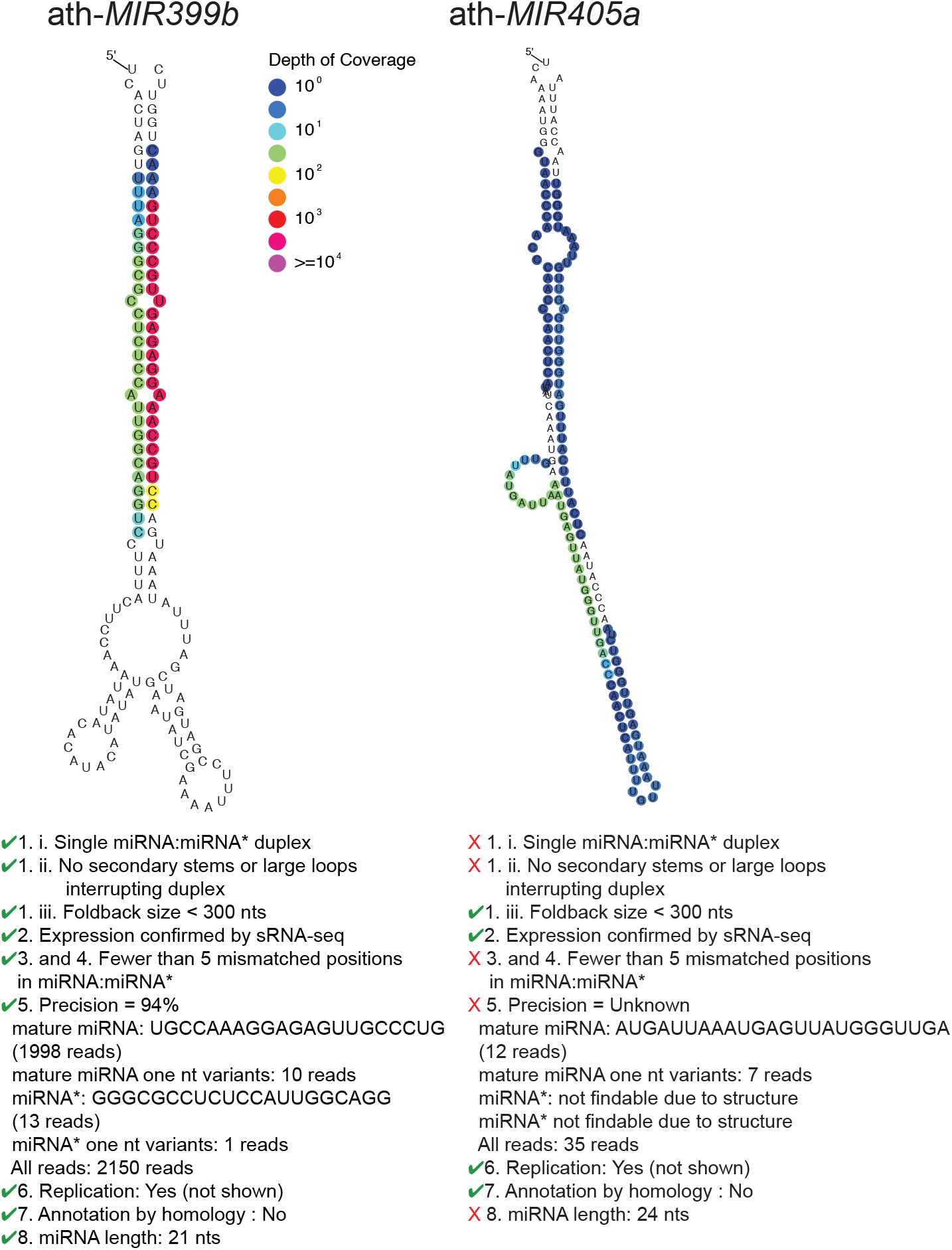
Examples of valid and invalid microRNA loci. Small RNA-seq data were from GSE105262 (Polydore and Axtell, 2017).Left: *Arabidopsis thaliana* (ath) *MIR399b*, a locus judged valid under the new criteria. Right: ath-*MIR405a*, a locus judged invalid under the new criteria.

Why is avoiding false-positive miRNA annotations of critical importance? The major issue is that, given the complexity of plant endogenous small RNA populations, even very small rates of false-positive miRNA annotations lead to overwhelming numbers of bad annotations. Many computational tools for plant miRNA discovery have been described, but they vary widely in performance. Lei and Sun (Lei and Sun, 2014) performed an informative benchmarking experiment in which the same small RNA datasets from *Arabidopsis thaliana* were used as input to seven different miRNA annotation tools (Figure 2). In these analyses, we assume for the sake of argument that the miRNA loci found by a program that did not already exist in miRBase (the central miRNA registry – see below) are false positives. Strikingly, several tools found more than 1,000 false positives during their analyses, and consequently have poor precision and accuracy. Lei and Sun’s analysis suggested that miR-PREFeR (Lei and Sun, 2014) and ShortStack (Johnson et al., 2016) performed the best. However, even the best tools had false positives, along with accuracies that rarely surpassed 60% (Figure 2). Plus, the software landscape is continually evolving so it is likely that these tools will improve or better ones may yet emerge. In light of progress in machine learning approaches, a possibility in the future of miRNA annotation is to efficiently sort small RNAs and classify miRNAs with a higher degree of accuracy than current methods, perhaps even without a reference genome. Indeed, the recent description of the Mirnovo tool (Vitsios et al., 2017) is promising, although its performance on plant datasets is diminished relative to animals. Whatever tool or internal procedure is used to annotate miRNAs from sRNA-seq data, users should exercise caution and common sense, and ask whether the procedure conforms to all of the annotation guidelines described above. In particular, any analysis that results in thousands of new miRNA annotations is quite likely to be comprised almost exclusively of false positives (Taylor et al., 2017), because even in intensively studied species like *Arabidopsis thaliana*, at most a few hundred miRNAs are known. Researchers, peer reviewers, and consumers of the published literature should be skeptical of a report that describes 100+ “new” miRNA loci in a given plant species.

**Figure 2.**
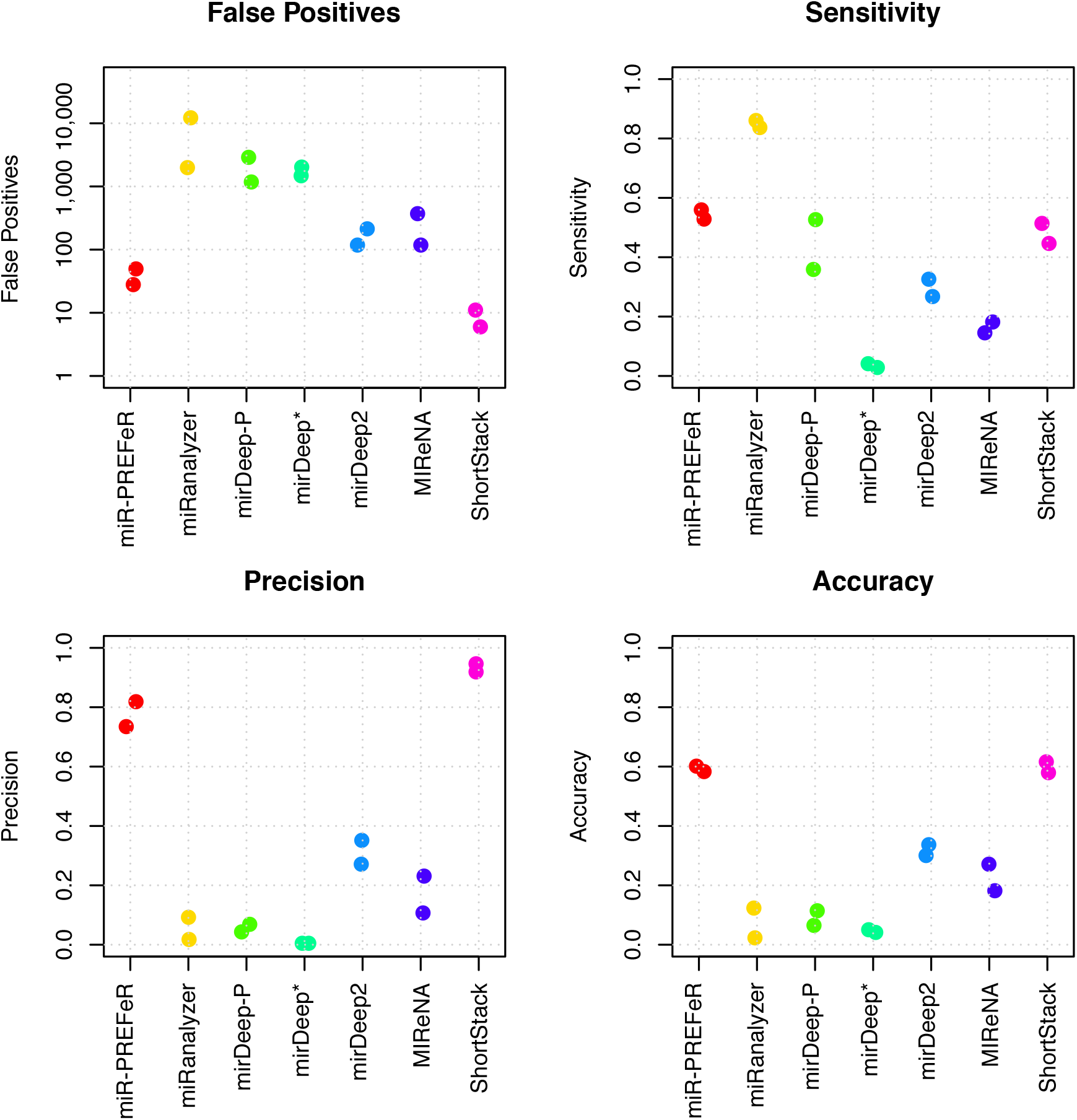
Comparative performance of miRNA annotation software from plants. Data are adapted from Lei and Sun (2014). In that publication, two *A. thaliana* sRNA-seq datasets were analyzed with seven different programs. miRNA loci that the programs found that were not in miRBase (version 20) were designated false positives. True positives: miRNA loci found that were also annotated in miRBase. False negatives: Expressed, miRBase-annotated miRNA loci not found by the tool. True negatives: miRBase-annotated miRNA loci that were not expressed and thus not found by any of the tools.

## The State of miRBase

miRBase (www.miRBase.org) is the centralized registry and database for miRNA annotations across all studied species (Kozomara and Griffiths-Jones, 2014). Its primary purposes are to assign consistent nomenclature to miRNA loci and to allow easy community access to all known miRNAs and their sequences. As of this writing, miRBase is at version 21, containing 6,942 miRNA hairpin annotations from 72 different land plant species; unfortunately, it has been over three years since this last release. miRNAs are placed into families based on the similarity of the mature miRNA sequences; individual families are assigned distinct numbers in the order of their discovery. Multiple loci from the same family can be present in a single genome, and families are conserved across plant species. As a result, the 6,942 hairpins represent 2,408 distinct miRNA families of land plants.

It is important to realize that miRBase is not the primary gatekeeper in terms of enforcing the quality of miRNA annotations. Instead, miRBase essentially collates and standardizes annotations from the peer-reviewed literature. The burden of quality control for miRNA annotations instead falls on researchers, peer reviewers, and editors. Because of the reliance on peer review and its inherently inconsistent results, and because of the large risk of false positive annotations (see above), the overall quality of miRBase is sub-optimal. Taylor et al. (2014) provided an important retrospective analysis of all of the land plant miRNA annotations present in miRBase version 20. Their analysis, based on sRNA-seq-based processing patterns, found that 1,993 out of the 6,172 (32%) land plant miRNA loci lacked convincing, supporting evidence. These dubious loci were disproportionately ‘singleton’ families; miRNA families represented by just a single locus in a single species. Thus, in their analysis, Taylor et al. (2014) marked as questionable 1,351 of the 1,802 (75%) of the land plant miRNA families in miRBase version 20. To the uninitiated, the idea that three out of four plant miRNA families are false annotations is probably a shock. Nonetheless, the criteria that Taylor et al. (2014) used are largely consistent with the best-practice methods we outline above (Table 1), and thus these estimates of the problems with miRBase annotations are likely close to the mark.

The curators of miRBase have recognized the growing issue of false positive miRNA annotations (Kozomara and Griffiths-Jones, 2014), and they have taken action. Beginning with the release of version 20, miRBase allows crowd-sourced comments on individual annotations, including the ability to ‘upvote’, ‘downvote’ and comment on annotations, and in some cases links to Wikipedia entries describing each locus. miRBase has also designated a subset of miRNA annotations as ‘high-confidence’. High-confidence loci are designated based on the pattern of read-alignments from reference-aligned sRNA-seq data. The criteria are similar in spirit to our recommended best practices (compare Tables 1 and 2). Strikingly, there aren’t that many high-confidence land plant miRNAs: Just 587 of the 6,942 (8.5%) of the land plant miRNA loci in miRBase 21 are marked as high confidence. If we designate a miRNA family as high confidence provided it has at least one high-confidence locus, we observe that just 227 of the 2,408 (9.4%) land plant miRNA families are “high confidence” as of miRBase 21. Clearly, available data support a conclusion that a great many miRBase entries, perhaps the majority, are questionable. Mere presence of an annotation in miRBase should absolutely not be taken to prove that a miRNA annotation is ‘real’, and instead, miRNA annotations must withstand the test of time and secondary analysis by independent groups and datasets. Importantly, miRBase also displays aligned small RNA-seq data for several species. When available, inspection of these data can help users decide for themselves the reliability of miRNA annotations.

Other retrospective efforts to re-evaluate miRBase miRNA annotations have focused on individual species, including rice (Jeong et al., 2011) and soybean (Arikit et al., 2014). In these cases, extensive manual curation coupled with computational analyses were used to assess prior annotations in light of deep sRNA-seq coverage. These efforts are clearly valuable, but do have limits. One of the issues is that miRBase is not designed for bulk removal of clearly erroneous annotations (although the idea of separating high-confidence annotations from the general population is promising). Another issue with the static-list approach is that the effort required is large, while the fast-moving pace of data accumulation can change the results quickly. New sRNA-seq data arrive in public archives almost daily. Thus, over time, initially questionable miRNA annotations might become better supported, while others that initially looked strong might be questioned. Given the volume of incoming sRNA-seq data, it is unreasonable to expect any one person or group (like the miRBase curators) to maintain a fully up-to-date database of all available aligned reads across all plant species, as well as making available what would be a constantly evolving set of evaluations.

Finally, a problem of growing concern is the large length of time between updates to miRBase. As of this writing (the last week of 2017), no updates to miRBase have been released in more than three years, although a social media announcement has promised a new update soon. During this time, many new miRNA submissions have been submitted to miRBase, and received official registry numbers, but none have yet been released by the database. This presents a serious problem in that it has become difficult to assert with confidence that ‘new’ miRNA annotations are truly new; it is not feasible to scour every single published paper on miRNAs and hunt through supplemental data tables to find all previously annotated miRNA sequences for all species. Perhaps part of the issue in maintaining miRBase is simply the volume of submissions and the workload required for manual curation of each newly-submitted list of miRNAs. Slow release of new annotations may also have the effect of discouraging submission of new sequences. We suggest that either miRBase or the community should consider alternatives to the current system. We advocate for the development of a fully-automated system that would permit uploads of candidates in a standardized format without requiring direct curator input. Quality control would be maintained by keeping the requirement that new miRNA annotations are part of an accepted or published peer-reviewed article, and by requiring that submitters’ names and contact information be publicly listed next to their annotations. The automated system could also be designed to enforce the modified requirements we outline above. In particular, it could require the underlying small RNA-seq alignments to be uploaded, followed by automated analysis of the prospective miRNA annotations. Users could receive a report on each locus, and with loci only accepted if they pass the analysis. Accepted annotations would at the submitter’s discretion become instantly available, or available upon publication of the associated study, with names automatically generated and registered. We also envision a system in which other users could evaluate and rank previous annotations based on the criteria above (e.g. Table 1). Users could influence rankings of individual miRNAs via submission of additional data or reports.

## Conserved Land Plant miRNAs

Understanding the challenges inherent in miRNA annotation requires some background on their patterns of conservation across plant genomes. In 2004, Floyd and Bowman (Floyd and Bowman, 2004) made the seminal observation that the predicted miRNA target sites for miR166, one of the first miRNAs discovered in *Arabidopsis thaliana*, were conserved in all major land plant lineages, including angiosperms, gymnosperms, ferns, lycopods, mosses, liverworts, and hornworts. A subsequent microarray study showed that several miRNA families first described in *A. thaliana* were detectable in ferns, gymnosperms, magnoliids, and monocots, with a few also detectable in a lycopod and a moss (Axtell and Bartel, 2005). The subsequent 10+ years have seen numerous studies that have applied sRNA-seq to the task of identifying miRNAs in diverse plant lineages. The key enabling technologies in these studies have been the development of highly parallel, next-generation sequencing methods, and the assembly of complete genome sequences for several key plant species. From an evolutionary perspective, the genome sequences of the basal angiosperm *Amborella trichopoda* (Amborella Genome Project, 2013), the lycopod *Selaginella moellendorffii* (Banks et al., 2011), and of the moss *Physcomitrella patens* (Rensing et al., 2008) have proven especially critical, because they represent key clades in land plant evolution that are generally under-sampled in terms of genomic resources.

Defining a ‘conserved’ miRNA is an inherently slippery concept. When applied to miRNAs, conservation is generally a shorthand for the accumulation of identical or near-identical (one or two nucleotide differences) mature miRNA sequences, although the miRNA* and some flanking hairpin sequence can also sometimes demonstrate conservation (Chorostecki et al., 2017). One of the key issues is at what taxonomic level the conservation is observed. For the sake of this discussion, we define a ‘conserved’ land plant miRNA family as one which has been annotated in miRBase 21 in at least two of the eight following major taxonomic divisions: Eudicots-Rosids, Eudicots-Asterids, Eudicots-Basal, Monocots, Basal Angiosperms, Gymnosperms, Lycopods, and Bryophytes. We further constrain the definition to minimize false positives by requiring that at least one of the annotations is designated ‘high-confidence’ in miRBase 21. Using these criteria, we see that 36 miRNA families are ‘conserved’ (Figure 3). These include nine families (miR156, miR160, miR166, miR171, miR319, miR390, miR477, miR529, and miR535) with high-confidence annotations in both *P. patens* and at least one angiosperm. These nine miRNAs most likely arose prior to the existence of the last common ancestor of all land plants and have been conserved in most or all diversified lineages.

**Figure 3.**
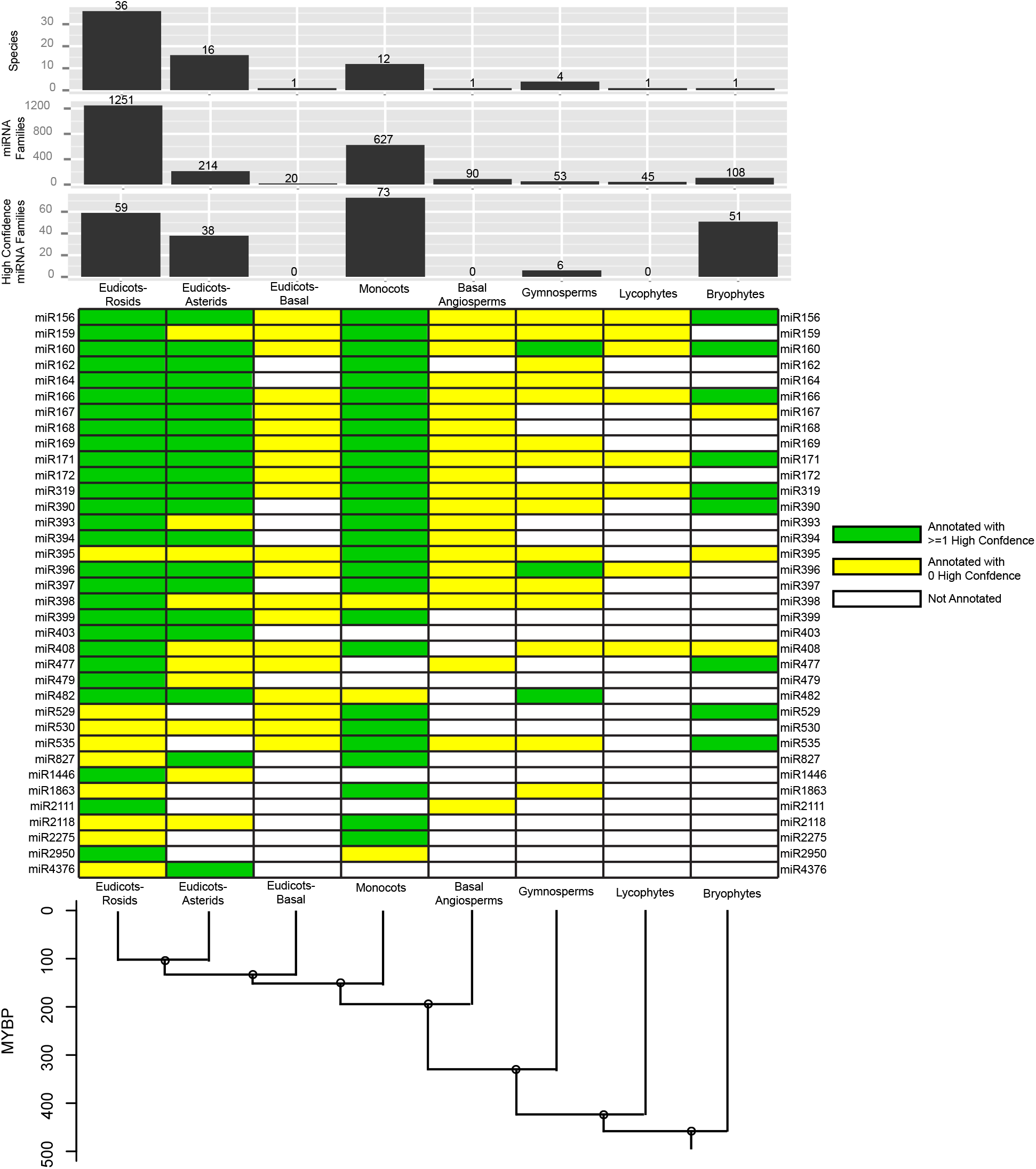
Conserved miRNA families in land plants based on miRBase 21 annotations. Annotations were binned into one of eight major taxonomic groupings. Top bar charts illustrate counts of the number of species, number of miRNA families, and number of high-confidence miRNA families in each group. The central heat map shows annotations for ‘conserved’ miRNA families, which are defined here as those annotated in at least two of the eight groups and which have at least one high-confidence annotation. The bottom cladogram illustrates the approximate divergence times (based on (Magallón et al., 2013) and (Wang et al., 2009)) of the eight groups. MYBP: Million Years Before Present.

Figure 3 also illustrates that there has been very uneven sampling density with respect to miRNA annotations in miRBase. In terms of species sampled and families annotated, the core eudicots (Rosids and Asterids), and monocots dominate miRBase. In contrast, basal eudicots, basal angiosperms, lycophytes, and bryophytes are each represented by just a single species (*Aquilegia caerulea, A. trichopoda, S. moellendorffii*, and *P. patens*, respectively). The uneven sampling density means that apparent patterns of loss in a lineage should be interpreted skeptically; they may well be artifacts of under-sampling. This is especially true in the basal eudicots, lycophytes, and gymnosperms. Indeed, a study of spruce small RNAs (Xia et al., 2015a) expanded the roster of conserved miRNA families in gymnosperms but is not yet reflected in the current version of miRBase, nor is a more recent study reporting extensive small RNA-seq data from many lycophyte and fern species (You et al., 2017). Therefore, there are still likely to be widely-conserved miRNAs that are not yet recognized as such, due to as-yet incomplete sampling of plant species and their small RNAs.

Proper miRNA annotations require both sRNA-seq data and the availability of a reference genome. This is because precursor hairpin structure is a key required feature needed for annotation, and while not essential, because the annotation of rRNAs and tRNAs can help to flag and remove their abundant decay products that can contaminate the output of miRNA prediction. This restricts *de novo* annotations of new miRNA families to species with sequenced genomes. Nonetheless, survey sRNA-seq data from species that lack a sequenced genome can still be useful to illuminate patterns of miRNA evolution and conservation. The disadvantages with this approach are that only those miRNA families that can be confidently annotated from ‘anchor’ species that do have reference genomes are countable. Cháves Montes et al. (Montes et al., 2014) reported a survey of land plant miRNA evolution based on an extensive sRNA-seq effort from 31 different vascular plant species, including one fern, three gymnosperms, and four basal angiosperms (but no bryophytes). Several patterns not apparent from miRBase annotations alone were identified. Perhaps most importantly, the observation of highly abundant miRNA reads from multiple species can buttress the annotation confidence for clades that are lightly sampled in miRBase. Specifically, the observation of high miRNA read counts from several miRNA families in multiple gymnosperm and basal angiosperm species (Montes et al., 2014) gives much more confidence in the existing miRBase annotations, which have few-to-no high-confidence annotations in miRBase at present (Figure 3). A large-scale, reference-genome-free, sRNA-seq effort from lycophytes and ferns was similarly able to buttress confidence of many more miRNA annotations (You et al., 2017).

## Mechanisms and Rates of Plant miRNA Emergence

As we demonstrate above and from numerous studies of miRNAs in early-diverged lineages, it is clear that individual miRNAs have emerged steadily and in parallel to the evolution of land plants. Publications describing the mechanisms by which plant miRNAs emerge are numerous, so rather than covering the topic in detail, we will simply note the consensus points. In miRBase version 21, there are 2,026 land-plant miRNA families (out of 2,408 total families) that are only annotated in a single species; of those, just 106 families are ‘high-confidence’; in other words, there are a variety of lineage-specific miRNAs plus a subset of miRNAs conserved to varying degrees. As we describe above, many apparently “species-specific” miRNAs may be misannotated siRNAs; ignoring those, there are still many non-conserved, lineage-specific miRNAs. Where do these come from and what is the process by which they emerge? Two primary mechanisms are clear: (1) spontaneous genomic formation via duplication of inverted repeats, and (2) gradual evolutionary shifts in miRNA sequences, in parallel with target gene divergence. In the first case, homology to target genes in miRNA precursors outside of the miRNA:miRNA* duplex supports inverted repeat formation as a step to miRNA generation (Allen et al., 2004); this is readily apparent among the highly redundant families that target nucleotide-binding site, leucine-rich repeat (*NLR*) ‘resistance genes’ (Zhang et al., 2016). The second case requires identification of sequence conservation across precursors of miRNAs, some of which may break the rules we describe above for designation of families. One example is the miR7122 “super-family” that includes miR173, miR7122, and miR1509 (among others) and shares a core sequence of just 13 nucleotides with the much more ancient miRNA, miR390 (Xia et al., 2013). The miR4376 super-family was a ‘stepping-stone’ to the identification of the emergence of miR7122, in that the miR7122-miR390 homology was weak and cryptic without knowing the intermediate homology of miR4376. The key lesson is that some miRNAs have ‘hidden conservation’, as yet unknown due to sampling bias; as new species’ genomes are examined that fill in the gaps between the limited number of sequenced plant genomes and characterized miRNAs, connections between “new” and “conserved” miRNAs are likely to be found. In the near term, this confounds efforts to estimate birth-and-death rates of plant miRNAs. Overall, while it is clear that many apparently lineage-specific miRNAs are probably erroneous annotations (Taylor et al., 2014), it also remains clear that many others are real. miRNAs without obvious conservation will continue to be a major part of future annotation efforts in plants.

## Conservation and Identification of miRNA Targets

For the most widely conserved plant miRNAs, their targets are also conserved (for a good list of these well-known conserved targets, see Table 1 from (Jones-Rhoades, 2012)). Indeed, purifying selection offers an easy explanation for conservation of miRNA sequences and their target sites: If a given miRNA-target interaction is critical for fitness, most mutations would be deleterious, as they would be most likely to decrease the complementarity between the miRNA and the target. This is especially true for miRNAs that have multiple important targets, as simultaneous mutations in the miRNA and all of the target sites that maintain base-pairing are highly unlikely.

Layered on top of this static continuity of old miRNAs and their old target sites is the observation that old miRNAs can pick up new, lineage-specific targets. miR396, which appears to have arisen before the last common ancestor of the seed plants (Figure 3) provides some striking examples. The ‘old’ miR396 targets are *Growth Regulating Factor* (*GRF*) mRNAs, which have been confirmed as targets in the eudicot *A. thaliana* (Jones-Rhoades and Bartel, 2004), the monocot *Oryza sativa* (Li et al., 2010), and predicted as targets in the lycopod S. *moellendorffii* (Debernardi et al., 2012). In the Brassicaceae and Cleomaceae (both in the Rosids supergroup of eudictos), miR396 also targets *Basic Helix-Loop-Helix 74* (*bHLH74*) homologs (Debernardi et al., 2012), while in sunflower (*Helianthus annus*, in the asterid supergroup of eudicots) it has gained a *WRKY* transcription factor target (Giacomelli et al., 2012). Gains of lineage-specific targets have also been described for the ancient miR156, miR159, miR167, miR398, miR408, and miR482 families (Zhai et al., 2011; Buxdorf et al., 2010; Chorostecki et al., 2012; Brousse et al., 2014; Xia et al., 2015b). These lineage-specific targets frequently have complementarity patterns that include bulged nucleotides, rendering them undetectable by standard plant miRNA target identification software. Alternative approaches that are more sensitive to these diverse complementarity sites have been described to meet this challenge (Chorostecki et al., 2012; Brousse et al., 2014).

Studying the conservation of plant miRNA targeting necessitates robust methods to conclusively identify targets in the first place. To the best of our knowledge, all functionally-verified miRNA-target interactions in plants have a high degree of miRNA-target complementarity (Wang et al., 2015). This is in contrast to animal miRNA-target interactions, which most frequently involve far less base-pairing (Bartel, 2009). Nonetheless, simplistic searches for plant miRNA targets with perfect or near-perfect complementarity are sub-optimal, because experimental data show there are clear position-specific effects; mismatches, G-U wobbles, and bulges have much stronger effects at some positions than others (Mallory et al., 2004; Schwab et al., 2005; Liu et al., 2014). Therefore, several methods for prediction of plant miRNA targets that take the position-specific effects into account have been described. These methods vary greatly in their ease of use, run-times, sensitivities, and false-positive rates (Srivastava et al., 2014). The sequences immediately flanking miRNA target sites can also affect target site efficacy (Li et al., 2014a; Fei et al., 2015; Zheng et al., 2017), likely because of local mRNA secondary structures. However, these effects are not yet confidently predictable from primary mRNA sequence and as such have not been reduced to practice in *de novo* miRNA target site predictions.

Regardless of how good the prediction method is, the output is essentially just a series of sequence alignments, which are not in and of themselves conclusive findings. We question the utility of publishing long lists of miRNA-mRNA alignments without any experimental validation, especially since the lists themselves are readily reproducible by any interested party and represent untested hypotheses, not empirical conclusions. Prediction lists comprising tens, hundreds, or more target sites for a single plant miRNA should be treated with extreme skepticism, as should meta-analyses of such lists (such as gene ontology [GO] enrichments). Such large numbers of targets are exceptional for a single plant miRNA, and there are strong theoretical and empirical arguments against the possibility of this occurring (Li et al., 2014b). If large numbers of targets for a single miRNA are found, there should be direct experimental evidence to support the assertions. Such exceptional cases do exist, such as the thousands of targets of the miR2118 family in rice that are experimentally supported by phasiRNA accumulation (Johnson et al., 2009). Those attempting to find plant miRNA targets should also be skeptical of inferring mechanistic attributes from the complementarity pattern. In particular, the psRNATarget server (Dai and Zhao, 2011) marks miRNA-target alignments containing central mismatches as leading to translational repression. This contradicts a substantial amount of empirical data. Many plant miRNA targets that are perfectly paired in the central region are nonetheless also translationally repressed (Chen, 2004; Yang et al., 2011; Li et al., 2013). Conversely, not all targets that do have central mismatches are translationally repressed: Some are non-coding RNAs that are still sliced despite the central mismatches (Allen et al., 2005), some act as target-mimics (Ivashuta et al., 2011; Liu et al., 2014), while others are simply non-functional, especially when the miRNA levels are not overwhelming (Li et al., 2014a).

## Conservation, Evolution, and Annotations of Endogenous siRNAs

As listed above, the two prominent, endogenous siRNA populations in plants are phasiRNAs and hc-siRNAs. Both present unique challenges to annotation both because their sheer abundance can lead to appreciable false positives when annotating miRNAs, and because of their differing patterns of conservation. PhasiRNAs are defined above, and are mentioned here because they may be misidentified as miRNAs, due to their 21- or 22-nt length, reproducibility across libraries and/or tissues, enrichment in specific tissues or cell types, and even their derivation from a processed inverted repeat. For example, rice miR5792 is derived from a locus that yields 24-nt phasiRNAs only in meiotic stage anthers (Fei et al., 2016). Therefore, separation of miRNA candidates from phasiRNAs requires careful, often manual assessment of each locus.

Endogenous hc-siRNAs comprise a major portion of the regulatory small RNA pool in key angiosperm model organisms including *A. thaliana* (Lu et al., 2006), *O. sativa* (Jeong et al., 2011), and *Zea mays* (Nobuta et al., 2008). Most hc-siRNAs are 24 nucleotides in length and are thought to function in RNA-directed DNA methylation, which couples 24-nt siRNA production from heterochromatic regions to targeting of chromatin-associated nascent non-coding RNA transcripts. This targeting, which can be (and perhaps often is) non-cell autonomous (Melnyk et al., 2011) results in the *de novo* deposition of 5-methyl cytosine at target loci (reviewed by (Matzke et al., 2015). In *A. thaliana*, complete removal of hc-siRNAs has only modest effects on genome-wide DNA methylation patterns (Stroud et al., 2013) and causes no obvious defects in organismal phenotype. In contrast, loss of hc-siRNAs in rice de-represses hundreds of protein-coding mRNAs, many of which are proximal to 24 nt-generating Miniature Inverted repeat Transposable Elements (MITEs) (Wei et al., 2014). These features are relevant in the context of miRNA annotation, because MITEs often contain miRNA-like inverted repeats, and heterochromatic regions may be low copy in the genome; hc-siRNAs from such regions may thus confound miRNA prediction algorithms and end up on output lists as candidate miRNAs. This is a particular challenge due to the somewhat circular argument that defining and annotating ‘heterochromatic’ regions in a newly-sequenced genome often depends on the presence of 24-nt siRNAs.

sRNA-seq samples from several non-angiosperm plants lack an immediately obvious signature of abundant 24-nt RNAs, including mosses (Axtell and Bartel, 2005; Arazi et al., 2005), lycophytes (Banks et al., 2011; You et al., 2017), conifers (Dolgosheina et al., 2008; Montes et al., 2014). These observations led to the suggestion that, unlike phasiRNAs and miRNAs, hc-siRNAs were not universal features found in all land plants (Dolgosheina et al., 2008). However, homologs of key genes known to be responsible for hc-siRNA biogenesis and function in angiosperms clearly exist in basal plant lineages (Zong et al., 2009; Banks et al., 2011; Huang et al., 2015; Wang and Ma, 2015; You et al., 2017). Reverse genetic analyses of these homologs, coupled with sRNA-seq analyses of the mutants, has shown that hc-siRNAs exist in the moss *P. patens* (Cho et al., 2008; Coruh et al., 2015), which implies that the pathway was most likely present in the last common ancestor of all land plants. Whether hc-siRNAs have been specifically lost in the conifers and/or ferns remains an open question, but the presence of high levels of 24 nt RNAs in specific tissues of the conifers Norway spruce (*Picea abies*) (Nystedt et al., 2013) and Japanese larch (*Larix leptolepis*) (Zhang et al., 2013) suggests that this may not be the case. Thus, as a class of endogenous plant small RNA, it is now clear that hc-siRNAs are probably as universally conserved in land plants as the miRNA and phasiRNA classes. However, there is no evidence that indicates the sequences of specific, individual hc-siRNAs are under the same level of strong, purifying selection as conserved miRNAs. Most hc-siRNAs arise from intergenic regions, especially from transposons and transposon fossils, and their presumed primary role is to recognize and suppress newer transposons, especially those near protein-coding regions (Zheng et al., 2012; Zhong et al., 2012). Thus, we’d expect any sequence conservation of individual hc-siRNAs to be restricted to conserved transposon domains, such as transposase or reverse transcriptase-derived regions. However, explicit studies of hc-siRNA sequence and locus-level conservation have not been reported, and this may be an interesting area for future study.

In contrast to miRNAs, there are not yet any community-accepted centralized databases dedicated to annotations of hc-siRNAs or phasiRNAs. This is remarkable considering their prevalence in most plant species. We propose that the plant sciences community would be well-served by a modernized, cross-species database that incorporates miRNA, hc-siRNA, and phasiRNA annotations. Centralization of data, along with a single registry of locus names, will greatly facilitate future research, especially for those who are not small RNA specialists.

## Prospects

The two key developments in the study of plant regulatory RNA diversity and evolution are genome sequencing and assembly projects from diverse species, and high-throughput sRNA-seq data. We expect that the future will see continued acceleration in new plant genome assemblies and in accumulation of sRNA-seq data. As in many areas of biology, the challenge has shifted from acquisition of data, to designing more robust analyses of the data. Increasing the rigor of small RNA annotations, especially of miRNAs and their targets, will be critical to prevent further degradation of central databases with overwhelming numbers of false positives. Increased use of transparent, easily reproducible sRNA-seq analytical methods should also be a key community goal to ensure that the deluge of small RNA data is put to good use.

**Table 2.** miRBase criteria for high-confidence miRNA annotations (reproduced from (Kozomara and Griffiths-Jones, 2014))

|  |  |
| --- | --- |
| 1 | At least ten sRNA-seq reads aligned with no mismatches to both the mature miRNA and the miRNA* |
| 2 | The most abundant reads from each arm of the hairpin precursor must pair in the miRNA/miRNA* duplex with 0 to 4 nt overhang at their 3' ends. |
| 3 | At least 50% of reads mapping to each arm of the hairpin precursor must have the same 5' end. |
| 4 | The predicted hairpin structure must have a folding free energy of < 0.2 kcal/mol/nt. |
| 5 | At least 60% of the bases in the miRNA and miRNA* must be paired in the predicted hairpin structure. |

## Acknowledgements

We thank Margaret Frank, Patricia Baldrich, and Matthew Endres for critical comments on this manuscript. We thank Seth Polydore for aligned sRNA-seq data. The US National Science Foundation supports research in both authors’ labs (MJA’s lab by award 1339207, and BCM’s lab by awards 1257869 and 1339229).

## Author Contributions

MJA and BCM wrote the manuscript. MJA analyzed data and prepared figures and tables.

